# Cloudgene 3: Transforming Nextflow Pipelines into Powerful Web Services

**DOI:** 10.1101/2024.10.27.620456

**Authors:** Lukas Forer, Sebastian Schönherr

## Abstract

**Summary:** Nextflow has emerged as a flexible and scalable workflow system for building computational pipelines in life sciences. Simultaneously, Software-as-a-Service platforms enable researchers to concentrate on data analysis rather than on setting up pipelines locally. However, converting existing pipelines into web services can be challenging due to complexities such as handling large datasets, web development and overall security. Cloudgene 3 addresses these challenges by providing a platform that transforms Nextflow pipelines into scalable web services in just a few steps. Making pipelines available as public or private services enhances collaboration among diverse teams and improves accessibility, eliminating the need for extensive technical expertise.

**Availability and Implementation:** Cloudgene 3 is free available at https://www.cloudgene.io

## 1 Introduction

The vast amount of biological data generated from various sources, including genomics, proteomics, and metabolomics, has transformed the life sciences into a computationally intensive field. Computational workflow systems like Nextflow have become game-changers for research institutes and companies worldwide, enabling them to analyze data in a reproducible and efficient manner (Di Tommaso, et al., 2017). Additionally, Software-as-a-Service (SaaS) tools in life sciences have gained popularity as they allow scientists to focus on data interpretation rather than the complexities of data analysis. By using these web tools, researchers can easily analyze their data, avoid common pitfalls, and streamline their research processes. Furthermore, SaaS facilitates access to datasets that may not be publicly available or cannot be shared broadly. For instance, the Michigan Imputation Server (Das, et al., 2016) allows users to impute genomes using a private reference panel through a web service. This provides a valuable resource to the community, as the reference dataset used cannot be shared and the imputation process is computationally intensive. Platforms like Michigan Imputation Server and mtDNA-Server (Weissensteiner, et al., 2024) both implemented as Nextflow pipelines, have gained widespread acceptance within the scientific community. These services highlight the value and effectiveness of web-based bioinformatics tools.

Nevertheless, transforming an existing Nextflow pipeline into a web service can be complex and time-consuming. It requires expertise in managing large datasets, web development and general security to protect sensitive information and data. Cloudgene 1 (Schönherr, et al., 2012) was designed to assist end-users in executing workflows using big data technologies like MapReduce. Unlike Galaxy (Galaxy, 2024), it was developed specifically for the Hadoop MapReduce environment. However, compared to widely used systems like Galaxy, Cloudgene 1 is not equipped to handle the demands of public web services that support numerous active users and thousands of jobs. In contrast, Seqera Platform (https://seqera.io) and Lifebit Copilot (https://www.lifebit.ai) enable an efficient management and execution of pipelines providing all available pipeline parameters across different cloud providers. However, their primary purpose is not to build use case-specific web services.

Here, we present Cloudgene 3, a platform for constructing scalable web services based on Nextflow pipelines. With features such as user management, a notification system, workflow administration, and monitoring of jobs, Cloudgene 3 provides an all-in-one solution without requiring web development knowledge for integration. Designed for easy usability, Cloudgene 3 allows users to upload files via its graphical user interface and start computational analyses instantly, eliminating the need to run pipelines locally. Existing Nextflow pipelines can be easily integrated and shared with the scientific community, enhancing collaboration in the field of life sciences. It is freely available and extensive documentation is provided on the website.

## 2 Methods

Cloudgene 3 is implemented in Java using the Micronaut 4 framework. It shares the same requirements as Nextflow and operates on all Linux systems. The software can be deployed with either a MySQL database or a self-contained H2 database, facilitating installations on cloud platforms such as AWS. Cloudgene 3 serves as a Platform as a Service (PaaS), offering end users SaaS functionality through the integrated Nextflow pipelines, referred to as Cloudgene apps. These apps are portable, self-contained units that are easy to share, install, and transfer between instances, requiring no modifications to the underlying pipeline. Each app includes a YAML file containing metadata, input/output parameters, and steps for chaining Nextflow pipelines. Applications can be hosted on GitHub and can be installed with a single command.

The framework integrates with the Nextflow ecosystem by utilizing its web-log functionality, which provides users with real-time feedback on running tasks. Cloudgene generates a unique secret URL with a random hash (collector URL), which is shared with Nextflow. Nextflow tasks send status updates to this URL whenever they start or finish. Cloudgene processes these events and updates the progress views accordingly. Once a task is complete, Cloudgene 3 checks for the presence of special report commands in the output of a task. This enables communication between the tasks and Cloudgene 3, allowing for error messages or status updates. Furthermore, Cloudgene 3 allows the configuration of Nextflow parameters and available executors through a comprehensive administration panel on an application basis, supporting execution engines such as Slurm or AWS Batch and cloud storage options like AWS S3.

All input data and temporary data are automatically deleted after execution, while the results are deleted after a configurable period. Cloudgene 3 also utilizes the Nextflow’s caching mechanism to ensure that completed tasks are not re-executed, preventing unnecessary resource consumption (see Figure 1). Moreover, Cloudgene 3 implements user authentication, role management and fine-grained permissions at the application level.

**Figure 1:**
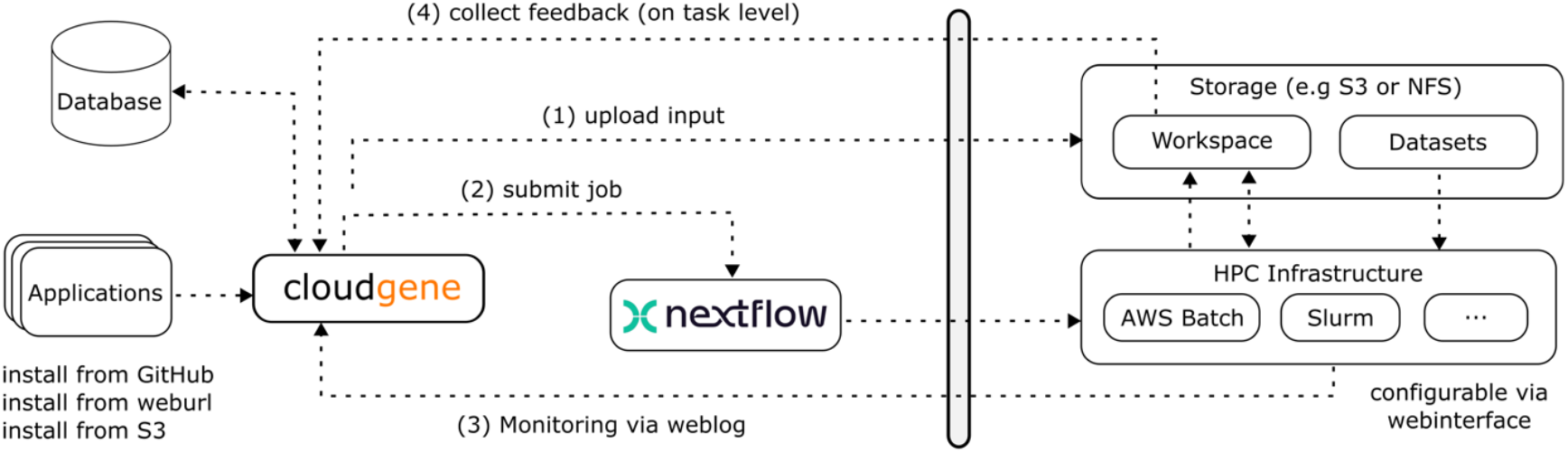
Integration of Nextflow and its communication within Cloudgene 3

## 3 Results

### 3.1 Pipelines as a Service

Turning a Nextflow pipeline like nf-core/fetchngs into a Cloudgene app involves a few key steps. First, the Nextflow pipeline must be connected to Cloudgene 3 by creating a configuration file that defines the inputs, parameters, and outputs for the pipeline. For nf-core/fetchngs, the app accepts a list of IDs as input, which the pipeline then uses to fetch sequencing data. An output parameter of type *folder* must be defined to store the downloaded data and the previously created sample sheet (see Figure 2). Once configured, the application can be installed on a Cloudgene 3 instance, which automatically generates a web interface based on input parameters, requiring no web development skills. This web interface can then be used by scientists to (a) upload their data and to set all provided parameters, (b) submit and monitor jobs and (c) download results or share them with collaborators through private links.

**Figure 2:**
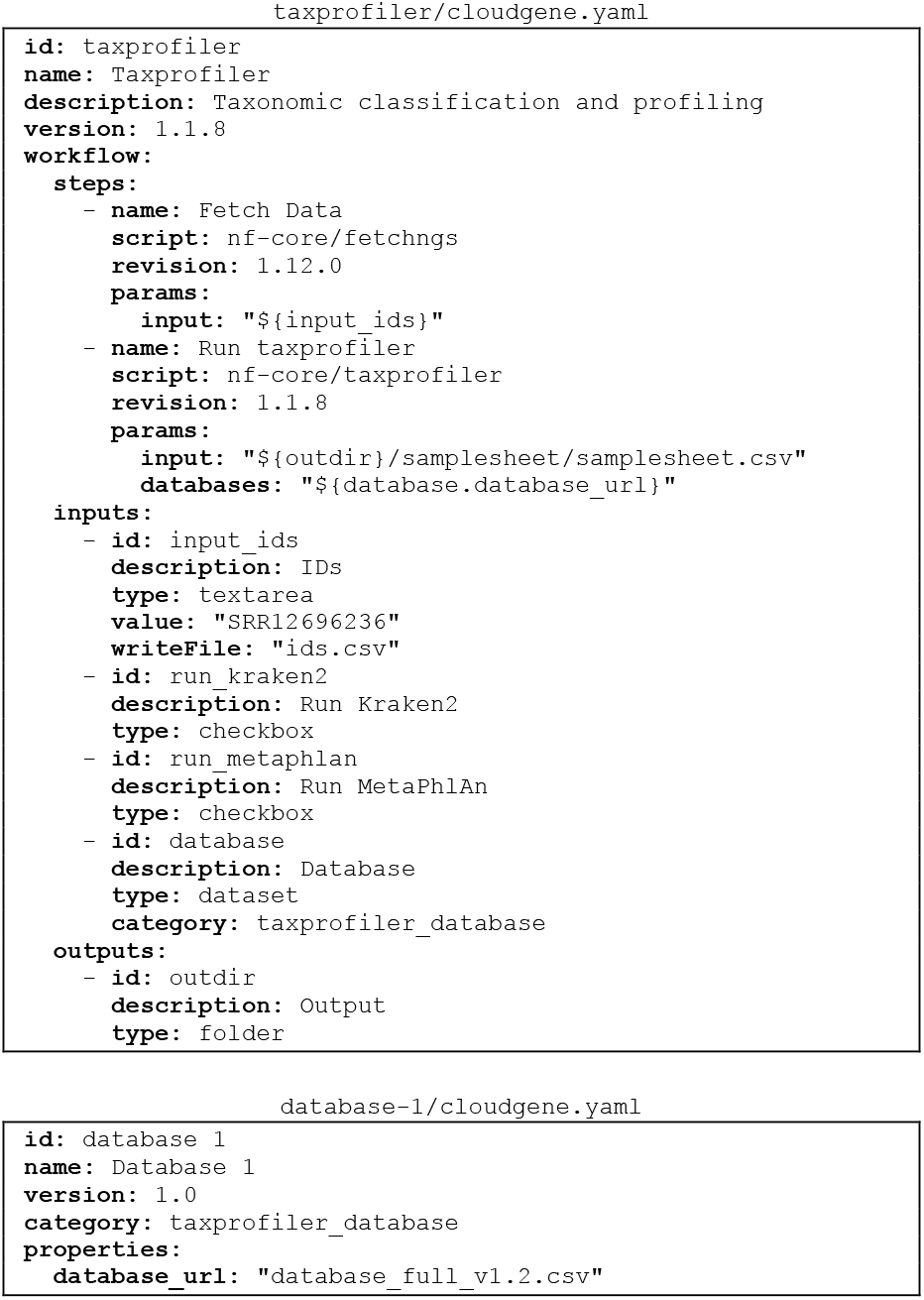
Example of a Cloudgene application that connect two pipelines and integrates datasets

### 3.2 Workflow Chaining

Cloudgene 3 allows the combination of multiple Nextflow pipelines by sequential steps. This means that the output of one pipeline can serve as the input for another, supporting complex data processing workflows. For instance, the sample sheet generated by the nf-core/fetchngs pipeline can be automatically mapped to the input parameters of a subsequent pipeline, such as the nf-core/taxprofiler (see Figure 2). This capability enhances the interoperability of different workflows and enables the development of more sophisticated applications by combining existing pipelines for specific use cases.

### 3.3 Decoupling Data from Pipelines

Datasets are conceptualized as applications without steps that can be integrated and linked with existing applications. The properties of a dataset can be used as input parameters of the pipeline, allowing users to choose one of the installed datasets from a given category (see Figure 3).

**Figure 3:**
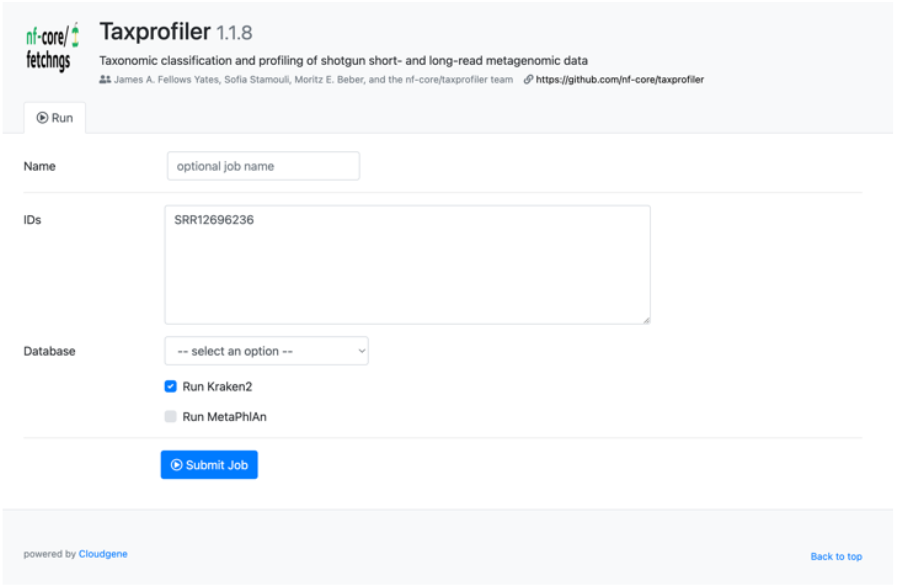
Example of the automatically created web interface for taxprofiler

Separating the dataset management logic from the pipeline itself has several advantages: First, dynamic dataset selection allows users to choose from a predefined list without modifying the pipeline configuration. Second, it promotes collaboration by allowing other users and institutions to integrate and share their datasets. Finally, permission management allows administrators to control access to specific datasets.

### 3.4 Existing Services

Cloudgene 3 serves as the backbone for several freely available large-scale web services in the field of genomics. The Michigan Imputation Server has processed over 116 million genomes and serves more than 12,000 users, facilitating high-quality genotype imputation for researchers world-wide (Forer, et al., 2024). The TOPMed Imputation Server (Taliun, et al., 2021) has imputed over 68 million genomes and supports over 5,000 users. Additionally, mtDNA-Server 2 provides a variant and heteroplasmy detection service for mitochondrial DNA and has analyzed thousands of samples (Weissensteiner, et al., 2024). Finally, haplocheck enables researchers to check mitochondrial genomic data for contamination (Weissensteiner, et al., 2021). These services demonstrate the usability, user-friendliness and stability of Cloudgene 3. Moreover, several nf-core pipelines (Ewels, et al., 2020; Langer, et al., 2024) have been encapsulated into web services and are provided on the website.

## 4 Conclusion

Cloudgene 3 represents a significant step toward establishing a standardized environment for deploying and using web services in the field of life sciences. The platform is fully self-contained, customizable and can be deployed on local infrastructure or in the cloud. Scientists can transform their Nextflow pipelines into powerful web services without the need to alter the pipeline itself, the deployment model, or the functionality. This integration opens new possibilities for sharing pipelines and deploying web services in a scalable and efficient manner.

## Acknowledgments

We would like to express our gratitude to all users of our web services for their valuable feedback. Special thanks go to Christian Fuchsberger, Albert V. Smith, Andrew P. Boughton and all GitHub contributors.

